# Global adaptation to climate change in the twilight zone revealed by shared signals of selection in mesopelagic lanternfishes

**DOI:** 10.64898/2026.05.22.727234

**Authors:** Bruna Cama, David Tian, Naomi Siu, Ben Frable, Ximena Prado, Malia Yalisove, Lydia Smith, Alexandra Dowlin, Sonke Johnsen, Anne Gro Vea Salvanes, Z. Jack Tseng, Adrienne Correa, Dahiana Arcila, Christopher H. Martin

## Abstract

Rapid accumulation of greenhouse gases threatens humanity and global diversity. The oceans absorb 30% of anthropogenic carbon emissions annually, but adaptation to climate change by the biotic components of this sink are poorly understood. Lanternfishes (Myctophiformes) are the most abundant vertebrates on the planet by biomass and the dominant mesopelagic vertebrate consumers, thus crucial components of the global carbon cycle. However, it is unknown whether lanternfishes are adapting to global warming and ocean acidification (OA). We hypothesized that warming and OA would act as major shared selective forces across diverse oceanic environments and that disparate taxa would respond in parallel through shared genetic pathways. We used whole-genome sequencing to test this hypothesis by identifying shared signals of selection across lanternfishes from multiple sites in the Atlantic and Pacific spanning three genera (*Benthosema glaciale, Triphoturus mexicanus,* and *Diaphus theta*). Across all species we found evidence of expansion from a population bottleneck possibly corresponding to the last glacial maximum and effective population sizes of only 5 million, suggesting substantial reproductive skew and spatially restricted populations. We successfully identified 34 candidate genes experiencing strong shared selection pressure across all taxa in both oceans. 81% of these candidate genes were consistent with adaptations to warming and OA, including a heat-shock protein (HSP70) and genes related to skeletal development, calcium homeostasis, and biomineralization. 14 out of 34 candidate genes are also known from experimental climate change studies to be involved in the response to hypoxia, altered pH, and thermal stress. We found significant gene ontology enrichment within these candidates for otolith morphogenesis, a major component of OA adaptation in fishes. This study provides a new approach for studying climate change adaptation at a global scale and our results imply widespread shared adaptive responses of marine species to climate change.

## Introduction

The oceans form the largest carbon reservoir on the planet, absorbing over 2 billion tons of carbon per year (DeVries et al., 2023; Nellemann & Corcoran, 2009; Venegas et al., 2023). Through the marine carbon pump, atmospheric carbon dioxide (CO2) is transported into the oceanic basins (NOAA, 2025). In the biological component of this marine carbon pump, CO2 enters the marine food web via phytoplankton and any organic carbon that is not remineralized below approximately 400m remains sequestered for hundreds of years or longer (DeVries et al., 2012; Nellemann & Corcoran, 2009; Nowicki et al., 2022). For example, sinking fecal pellets from krill in the Southern Ocean alone annually sequester an equivalent amount of carbon to all coastal ecosystems combined (Cavan et al., 2024).

Rising atmospheric CO2 levels are causing increasing ocean acidification (OA), rising water temperatures with more frequent and longer lasting heatwaves (Masanja et al., 2023; Oliver et al., 2018), and decreased oxygen availability and delivery (Frölicher et al., 2018; Gobler & Baumann, 2016; Hofmann & Schellnhuber, 2009; Oliver et al., 2018; Pörtner, 2008), with higher incidence and volume of hypoxic zones (Altieri & Gedan, 2015; Hofmann & Schellnhuber, 2009; Sampaio et al., 2021) and increased blood cell mortality (Meseck et al., 2016; Srinivasan et al., 2022; Wu et al., 2016). The effects of these environmental changes have been documented primarily in marine invertebrates: for example, by reducing calcium availability, OA impairs the development of mineralized tissues in corals (Armstrong & Bahr, 2025), foraminifera (Kuroyanagi et al., 2021), and shelled molluscs (Duquette et al., 2017; Fitzer et al., 2018); in bivalves, hypoxia affects growth rate, survival, and shell growth in complex ways, triggering altered respiration rates (Stevens & Gobler, 2018).

Despite potential ecosystem-wide effects of ocean acidification, there are few studies of its impacts on any marine fish, and most of these studies were carried out only in the laboratory. For example, increased levels of dissolved CO2 result in reduced growth and severe tissue damage in Atlantic herring (Frommel et al., 2014) and reduced red blood cell count and anaemia in Asian sea bass (Srinivasan et al., 2022). Mineralization of developing skate embryos is affected by pH, with contrasting effects depending on temperature and body region (Di Santo, 2019). Additional work has consistently demonstrated detectable responses in fish bone density to OA challenges in the laboratory (Leung et al., 2022; Sundin, 2023). OA may also impact fish sensory systems in myriad ways (Ashur et al., 2017), though evidence of this is lacking (Clements et al., 2022). Thus far, OA research has focused on coastal fisheries that are amenable to laboratory experiments (Bernal et al., 2022; Devergne et al., 2023; Kempf et al., 2024; Swank et al., 2025). The lack of knowledge for non-model, non-commercial, and non-coastal fish species constitutes a severe blind spot, especially considering the disproportionate involvement of mesopelagic fish populations in the global carbon cycle (Davison et al., 2013; Pinti et al., 2023; Saba et al., 2021).

Most carbon that is transported from the surface to the ocean floor through the biological pump will eventually pass through the mesopelagic ‘twilight’ zone. Located 200 – 1000m below sea level, this region separates the sunlit epipelagic zone from the dark bathypelagic zone. Lanternfishes (Myctophiformes; approximately 250 species) are the dominant vertebrate inhabitants of the mesopelagic zone in all oceans except the Arctic, comprising over 40% of total mesopelagic fish biomass, and are the most abundant vertebrates on Earth by biomass (Catul et al., 2011; Gjөsæter & Kawaguchi, 1980; Irigoien et al., 2014). They are central mediators in the marine food web, serving as prey for a wide range of predators, while preying on zooplankton primary consumers (Catul et al., 2011; Choy et al., 2012). Lanternfishes also play a major role in the global carbon cycle, serving as both a sink and source of carbon due to their diel migration to surface waters (a behaviour shared across 70% of all myctophiforms) (Catul et al., 2011; Knutsen et al., 2023) and central role in carbon transport between the epipelagic zone and deep ocean (Cavan et al., 2024; Eduardo et al., 2021; Knutsen et al., 2023; Koz, 1995). Like all marine fishes, they also produce calcium carbonate within their guts and excrete this in surface waters, resulting in the net transfer of dissolved calcium bicarbonate (alkalinity) to the surface and counteracting increasing concentrations of dissolved CO2 in surface waters (Saba et al., 2021; Wilson et al., 2009).

Despite the outsized importance of Myctophiformes in the marine food web, global carbon cycling, and ocean biochemistry, nothing is known about their responses to climate change. Studies of natural selection, adaptation, or genomic signals of selection have not been conducted on any mesopelagic fishes. In this study, we used population genetics tools to detect recent selective sweeps in lanternfish taxa across Atlantic and Pacific oceans. This dataset includes four populations of lanternfishes spanning three species, three subfamilies (Lampanyctinae, Diaphinae and Myctophinae, all in the family Myctophidae), and four geographic locations: the East Bering Sea (Alaska, US), the Eastern Pacific (San Diego, California, US), the Western Atlantic (Baltimore Canyon, Maryland, US), and the Northern Atlantic (Fensfjorden, Norway). We predicted that global climate change may create similar selective pressures across distant Ocean basins and disparate taxa, potentially resulting in parallel selection on the same genes across Myctophiformes. We focus on signals shared across the same genes in all four populations with particular attention *a priori* to the following functional categories most directly relevant to climate change adaptation: A) bone development and calcification (biomineralization) (Di Santo, 2019); B) respiration and circulatory system (hypoxia); C) cellular damage in response to thermal stress (heat shock) (Hu et al., 2022); D) cellular damage in response to ocean acidification (acid stress); and E) calcium homeostasis. Thus, we gained insight into global selection pressures on lanternfish populations at an unprecedented scale, spanning the Pacific and Atlantic and both inner seas and open ocean, demonstrating a new approach for detecting adaptations to climate change across taxa and environments.

## Methods

### Specimen collection

Specimens collected for these analyses represent three major myctophid subfamilies (Fig. 1): Lampanyctinae, Diaphinae, and Myctophinae. Lampanyctinae is represented by *Triphoturus mexicanus* (n=20) collected from San Diego, CA, US in 2024. Diaphinae is represented by *Diaphus theta* (7 individuals) collected from the eastern Pacific (both East Bering Sea, AK, US [n=5] and San Diego, CA, US [*n* = 2]) in 2024. Myctophinae is represented by two *Benthosema glaciale* populations connected to the Atlantic Ocean: Baltimore Canyon off the coast of Maryland, US (*n* = 14) collected in 2018 and the inner waters of Fensfjorden, Norway (*n* = 16) collected in 2022 (Fig. 1B). All specimens were collected via midwater trawling, e.g. the Isaacs-Kidd Midwater Trawl on the R/V *Gordon Sproul* at the Scripps Institution of Oceanography and the Harstad trawl on the R/V *G. O. Sars* in the Norwegian fjords. Specimens were euthanized in a buffered solution of MS-222, preserved in 95-100% ethanol, and catalogued in the Museum of Vertebrate Zoology Fishes Collection (MVZ:Fish:879-882). See Supp. Table 1 for specimen details and coordinates.

**Figure 1.**
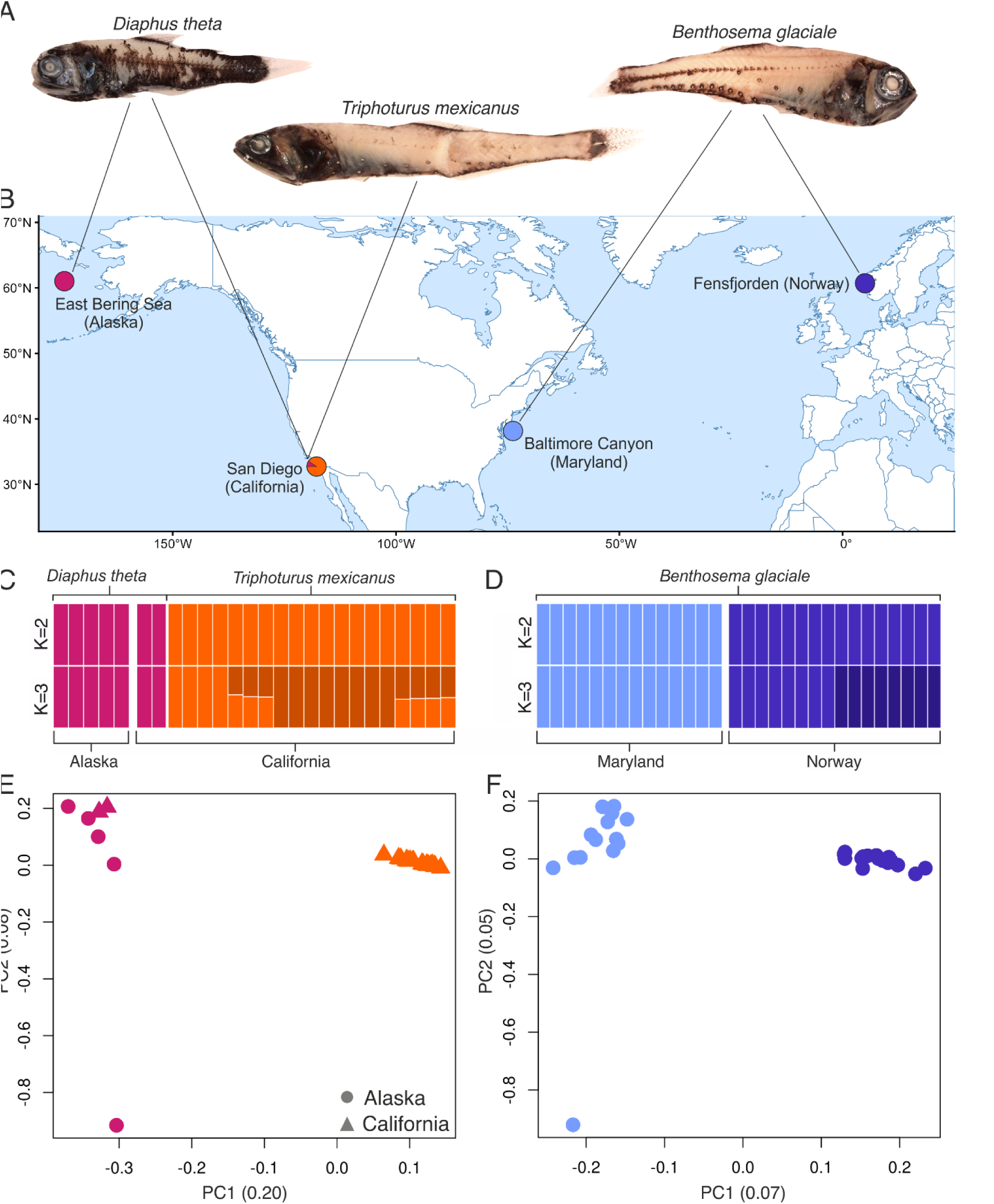
*A.* Photographs of the three focal Myctophid species, from left to right: *Diaphus theta, Triphoturus mexicanus,* and *Benthosema glaciale. B.* Northern hemisphere map showing the four collection sites, each connected to the species that were sampled at that location, with colors corresponding to representative species at that site. *C-D.* Admixture plots for K=2 and K=3. *E-F.* Principal component axes for LD-pruned whole genome data.

**Table 1.** Median π and Tajima’s D across *B. glaciale, T. mexicanus* and *D. theta,* with IQR reported in brackets. Values of π are shown averaged across the genome, across likely selective sweeps (CLR peaks output by SweepFinder2 above the 2% quantile threshold) (Fig. 3). and across regions not implicated in selective sweeps (CLR below the 2% quantile threshold). Tajima’s D is averaged across the genome.

| Species/region | Median $\pi$ (Nucleotide diversity) | | | Median Tajima's D |
| --- | --- | --- | --- | --- |
|  | Overall | In peaks | Outside peaks |  |
| *Benthosema glaciale*<br>(Norway) | 0.0012<br>(0.0005–0.0021) | 0.0008<br>(0.0003–0.0014) | 0.0012<br>(0.0005–0.0021) | -3.80<br>(-4.57 – -3.35) |
| *Benthosema glaciale*<br>(Maryland) | 0.0015<br>(0.0007–0.0026) | 0.0008<br>(0.0003–0.0013) | 0.0016<br>(0.0008–0.0027) | -3.90<br>(-4.60 – -3.43) |
| *Triphoturus mexicanus* | 0.0023<br>(0.0014–0.0035) | 0.0008<br>(0.00003–0.0023) | 0.0023<br>(0.0014–0.0035) | -2.99<br>(-3.48 – -2.42) |
| *Diaphus theta* | 0.0018<br>(0.0008–0.003) | 0.0006<br>(0–0.002) | 0.0018<br>(0.0008–0.003) | -5.40<br>(-6.36 – -4.52) |

### DNA extraction and whole-genome sequencing

DNA was extracted from fin-clips from each specimen with Qiagen blood and tissue kits and quantified with a Qubit 2.0 fluorometer to standardize concentrations before whole-genome library preparation using a modified Kapa Hyper Prep protocol (Benham et al., 2025) at the University of California, Berkeley. Genomic libraries were sequenced at the Vincent J. Coates Genomic Sequencing lab (QB3 Genomics, UC Berkeley, Berkeley, CA, RRID:SCR_022170) on a NovaSeq X 25B flowcell using 150 bp paired-end sequencing.

### Genomic sequencing

Read quality was assessed using a combination of fastQC (Andrews, 2015) and Qualimap 2 (Okonechnikov et al., 2016). Read mapping was carried out via a pipeline that employs Burrows-Wheeler Alignment (H. Li & Durbin, 2009) using the bwa-mem2 function, part of the bwa command-line software (H. Li, 2013), followed by sorting and duplicate removal using the open source package SAMtools (Danecek et al., 2021) and Picard toolkit (Broad Institute, 2019), respectively. The whole pipeline was run within a wrapper python script (De-Kayne et al., 2025), available on GitHub (DeKayne & Martin, 2025).

*Benthosema glaciale* samples were aligned to the fully annotated *Benthosema pterotum* reference genome (Q. Liu et al., 2024), while Lampanyctinae and Diaphinae samples were aligned to the partially annotated *Lampanyctus achirus* (formerly *Nannobrachium achirus* as it is catalogued on the Vertebrate Genome Project/NCBI) reference genome (Sanger Institute, 2022) (Supp. Table 1). Samples that aligned poorly to either reference genome (proportion mapped <50%) were considered misidentified or degraded and were removed from downstream analyses (n=2) with subsequent re-coding of allele frequencies. *L. achirus* was chosen as the most closely related reference genome for *D. theta* specimens based on the best mapping statistics obtained among all available lanternfish genomes at the time (mean proportion mapped approximately 70%). Unplaced scaffolds (<5000 bp, 554 for *B. pterotum* and 648 for *L. achirus*) in both reference genomes were not included in any downstream analyses due to poor mapping quality and to their unknown genomic location making them uninformative for sweep analyses.

Genotypes were called using BCFtools mpileup (Danecek et al., 2021) set to ignore indels with a maximum read depth of 250 per position per input file. Samples were aggregated into two separate VCFs based on their reference genome: one for all *Benthosema glaciale* samples and a Lampanyctinae/Diaphinae VCF for *Triphoturus mexicanus* and *Diaphus theta*. These two VCFs were filtered for genome quality (GQ≥30), read depth (DP≥3), and minor allele count (MAC≥1) using bcftools.

### Demographic history and population structure

To investigate the demographic history of myctophid populations, we estimated effective population sizes (N_e_) in GONe (Santiago et al., 2025) with input .PED/.MAP files generated from the *Benthosema glaciale* VCFs for Norway and Maryland and the *Triphoturus mexicanus* VCFs in PLINK 1.9 (Purcell et al., 2007) for the past 200 generations, corresponding to 100 years based on the myctophid generation time of six months (Sassa, 2019; Sassa et al., 2014). For this analysis only sites that were genotyped for over 80% of individuals were included, and *Diaphus theta* was fully excluded due to insufficient sample size. The recombination rate of Myctophidae is unknown, so we used three different recombination rates: 1.6 cM/Mb, the mean genome-wide recombination rate in zebrafish (Bradley et al., 2011); 2.54 cM/Mb from Atlantic herring (Pettersson et al., 2019); and 3.11 from threespine stickleback (H. Wang et al., 2026). We also estimated Tajima’s D and π (nucleotide diversity) in pixy (Korunes & Samuk, 2021) for all four populations. Log_10_(π) was computed for plotting, with a constant (10^-6^) added to all values to account for zeros during logarithmic transformation. This constant was chosen based on the smallest detected non-zero π.

To assess population structure, we used PCA in PLINK with linkage-pruned SNPs (*r^2^*, window size = 50 kb, step size= 5 kb), separately for Lampanyctinae/Diaphinae and Myctophinae. Only sites genotyped in over 80% of individuals were included in the PCA. The *Benthosema* dataset split into two subpopulations representing Baltimore Canyon (Maryland, US) and Fensfjorden (Norway) *B. glaciale.* The Lampanyctinae/Diaphinae clades were more difficult to differentiate as many individuals originally lacked genus-level identifications, so putative single-species clusters were identified on the pruned dataset via IBS (identity-by-state) in PLINK followed by k-means clustering analyses in R. Robustness was evaluated using the silhouette function in R(cluster). The most robust clusters recovered by this method were considered a single population (Supp. Fig. 1). One cluster corresponded to samples identified as *T. mexicanus* from San Diego, California and the other corresponded to samples identified as *Diaphus theta* from the East Bering Sea, Alaska and additional individuals from San Diego. Allele frequencies were updated with BCFtools following separation into the four subpopulations. Admixture between the two populations in each pair was assessed via PLINK on linkage-pruned SNPs with max k=4 (Fig. 1C).

For *T. mexicanus* and *Diaphus theta,* certain biallelic sites were marked as triallelic as the reference allele from *Lampanyctus ritteri* was never present in the genotyped individuals, due to their more distant phylogenetic relationship . These sites were recoded based on the frequency of the two alleles: the most frequent allele was recoded as 0 (major) and the least frequent as 1 (minor). This was appropriate because downstream scans for recent selection did not assume either of the two alleles was ancestral/derived.

### Genomic scans for selection

The four populations (Maryland and Norway *B. glaciale, Triphoturus mexicanus,* and *Diaphus theta*) were independently scanned for selective sweeps using SweepFinder2 (Degiorgio et al., 2016), one chromosome at a time, based on the folded site-frequency-spectrum (SFS). The hard selective sweep analysis was restricted to 100% genotyped biallelic sites and used a fixed grid size of 1000 bp. To focus on only the candidate loci showing the strongest evidence of selection, we searched across peaks whose composite likelihood ratio (CLR) was in the top 2%. This conservative threshold was chosen to capture only the strongest selective sweeps relative to the commonly used 5% threshold (Huang et al., 2022; McGirr et al., 2017; Z. Zhu et al., 2020). 2% was favoured instead of the more stringent 1% to account for potential differences in signal intensity among the three species due to differing sample sizes and coverage. Sweeps were compared between Maryland-Norway *B. glaciale* and between *Triphoturus-Diaphus* via positional alignment of results in R. Peak margins were calculated by detecting contiguous above-threshold regions using the rle (run-length encoding) function in base R. Any positional overlap was considered an indication of a shared selective sweep. 10 kb at the start and end of each chromosome was excluded from the search for shared peaks to account for repetitive elements located in the telomeres. To further assess recent selection, π (nucleotide diversity) was calculated separately for regions under the peaks and outside the peaks to test whether increases in the composite likelihood ratio for a sweep (CLR statistic) corresponded with local minima in nucleotide diversity.

We then extracted all genes from the reference genome annotations located within 50 kb of the CLR peak margins, conservatively spanning the range of distances between coding sequences and *cis-*regulatory regions based on existing data for fishes (Clément et al., 2020; Fang et al., 2020; Lee et al., 2015), to identify a set of candidate genes putatively under selection.

### Shared ontology and identity of shared genes

Candidate genes under positive selection were compared across the four populations at two different levels: shared genes across all four datasets and overlap in enriched gene ontology (GO) terms between GO analyses within each of the two oceanic basins. Gene identity was chosen over positional information due to substantial changes in synteny between the two reference genomes.

For both Atlantic and Pacific datasets, we used BLAST to compare the protein sequences of 2-way shared genes against the Swissprot database (Bateman et al., 2025). The BLASTp search was taxonomically restricted to *Danio rerio* (zebrafish), excluding all matches with E-values > 10^-6^. Once 2-way shared sweeps and the corresponding zebrafish genes were extracted from the Maryland-Norway *B. glaciale* and the *Triphoturus-Diaphus* comparisons, the two lists of candidate genes were pasted into UniProt search to obtain the corresponding zebrafish gene symbols and protein identities, later cross-verified on ZFIN. To test for shared gene ontology patterns between candidate genes in both datasets, R(clusterProfiler) (Yu et al., 2012) was used to assign ontology terms to each of these genes based on the zebrafish background and to test for GO enrichment with a *p-*value cutoff of 0.01.

Shared identity between recently selected genes across all oceanic basins was then determined by matching UniProt accessions/gene symbols between the two lists of 2-way shared candidate genes. Those found in both datasets were then extracted as shared candidate genes within or near CLR peaks in all four populations.

Due to the high percentage (∼73%) of *Triphoturus/Diaphus* sequences that could not be matched to any zebrafish gene, additional signals of homology among selected genes were discovered by performing BLASTp reciprocally between *Benthosema* and *Triphoturus/Diaphus* sequences under selection, excluding E-values > 10^-5^. Most genes in the *B. pterotum* genome are annotated with KEGG identities, which were attributed to the matching unidentified *Triphoturus/Diaphus* genes. This recovered 85 additional cases of shared homology between genes that had not been previously identified from zebrafish sequences. The zebrafish orthologs of these gene IDs were recovered and their ontology and identity were extracted from a combination of UniProt, KEGG (Kanehisa, 2000), and ZFIN databases (Bradford et al., 2022). When both KEGG and Swissprot BLAST identities were available for the same gene, the latter was favoured over the KEGG annotation due to its highly curated gene functional information. For genes lacking zebrafish homology, functional information was found through human, mouse or rat orthologs.

Because the number of 4-way shared genes was small (*n* = 34), unlike 2-way shared genes, an analysis of gene ontology enrichment for these candidates was not feasible; however, we were able to obtain individual ontologies from UniProt for all 4-way shared genes except 3 (*adam12a, clip, harbi1*) (Supp. Table 2). We defined six broad functions relevant to OA/climate change adaptation: thermal stress response, acid stress (high CO_2_ concentrations) response, hypoxia response, biomineralization, Ca^2+^ binding/modulation, and otolith formation. For each of the 34 genes, information was found in the literature and in ZFIN (Bradford et al., 2022) about their functional roles, with particular attention to past experimental climate change studies (Table 2). All genes with a documented function in the circulatory system were marked as likely relevant to hypoxia due to the circulatory system’s role in oxygen transport even when published experimental evidence of their expression patterns during hypoxia or in the context of OA could not be found. Likewise, all genes involved in skeletal system development were marked as relevant to biomineralization, and all genes involved in protein ubiquitination were marked as relevant to thermal stress response.

## Results

### Population structure

We found clear, consistent patterns of population structure using PCA and k-means clustering analysis. In the Atlantic, 14 *B. glaciale* from Baltimore Canyon (MD, US) were distinct from 16 conspecifics collected in Fensfjorden (Norway). In the Pacific, 19 *Triphoturus mexicanus* individuals from San Diego (CA, US) formed a single uniform cluster and the second-best supported cluster included seven individuals: two from San Diego and five from the East Bering Sea (AK, US), identified as *Diaphus theta.* Despite the heterogenous geographic origins of the *D. theta* cluster, genetic similarity within the cluster was high: its members shared a greater proportion of alleles than individuals within the *T. mexicanus* cluster. *T. mexicanus* were still unambiguously recovered as a single cluster, with strikingly consistent proportions of shared alleles between all individuals (Supp. Fig. 1). These results are reflected in both the PCA and the admixture plots (Fig. 1C-F).

### Demographic history

Using GONe, we estimated a stable effective population size of approximately 5 million for *T. mexicanus* and for *B. glaciale* in both Maryland and Norway over the past 200 generations (100 years) (Supp. Fig. 2). These results were consistent across different input recombination rates. Minor decreases in N_e_ are detected across species (20,000-40,000), but are negligible given our high esitmated N_e_ (Gargiulo et al., 2024). Genome-wide Tajima’s D was strongly negative across all four populations, consistent with recent population expansion (Table 1). Mean genome-wide π (nucleotide diversity) was 0.17% across the four populations (Table 1).

**Figure 2.**
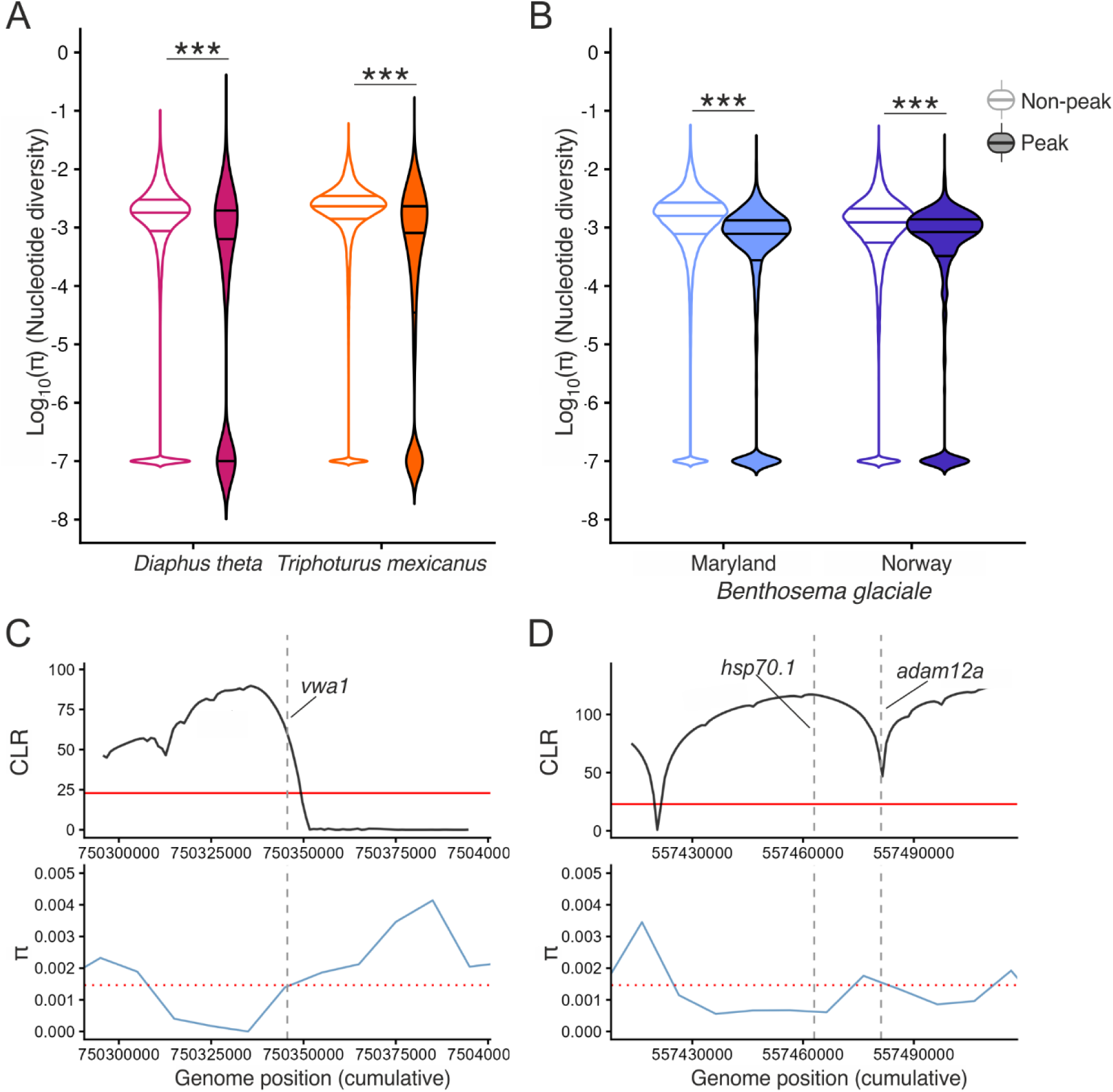
Reduced genetic diversity within hard selective sweeps. *A-B.* Reduced nucleotide diversity in peaks (1kb windows above the upper 2% CLR threshold) vs. non-peaks (1kb windows below the upper 2% CLR threshold) in the four focal populations. Horizontal bars within violins show the median and the 25%, 75% quantiles. *C-D.* Examples of genes located within selective sweeps showing increased compositive likelihood ratio (CLR) and decreased genetic diversity (π) relative to flanking regions in Maryland *B. glaciale*. Solid horizonal red lines in CLR plots show the 2% CLR threshold. Dotted horizontal red lines in π plot show the median for the species. Dashed vertical lines show the location of shared genes (Table 2). Cumulative genomic position is shown in bp. *C. Vwa1,* encoding an ossification/cartilage protein, occurs within a selective sweep. (Vargas et al., 2022; Yuan et al., 2025; T. Zhu et al., 2023). *D*. *Hsp70.1,* encoding a key heat shock protein implicated in experimental studies of climate change, and *adam12a* both occur within a selective sweep. All three genes shown in *C-D* occur in the proximity of selective sweep signals across all focal taxa (Fig. 3, Table 2).

### Shared selective sweeps across species and oceans

We successfully detected recent hard selective sweeps in both Atlantic and Pacific lanternfishes. To focus on only the candidate loci showing the strongest evidence of selection, we conservatively searched across peaks whose composite likelihood ratio (CLR) was in the top 2% of all peaks. In the Atlantic, we detected 396 peaks exceeding this threshold in Maryland *B. glaciale* and 429 peaks in Norway *B. glaciale.* In the Pacific, we detected 1,141 peaks exceeding this threshold in *T. mexicanus* and 531 in *Diaphus theta*.

Consistent with expectations, we found that these peaks generally corresponded to local dips in nucleotide diversity (Fig. 2): π within windows corresponding to CLR peaks (regions above the 2% CLR threshold) was 68% lower than π outside CLR peaks (Wilcoxon rank sum test, *P* < 0.00001) in Norway *B. glaciale* and 49% lower than π outside of CLR peaks (*P* < 0.00001) in Maryland *B. glaciale*. Likewise, π within peaks was 35% lower than π outside of CLR peaks in both *T. mexicanus* (Wilcoxon rank sum test, *P* < 0.00001) and *D. theta* (*P* < 0.00001) (Fig. 2). 94 peaks were shared between the two *B. glaciale* populations (Fig. 3A-B). Based on the existing *B. pterotum* annotation (Q. Liu et al., 2024), shared sweeps and their 50kb flanking regions spanned 523 coding regions (Fig. 3). 73 peaks were shared between *T. mexicanus* and *Diaphus theta* (Fig. 3C-D). These shared sweeps and their 50 kb flanking regions spanned 543 coding regions.

**Figure 3.**
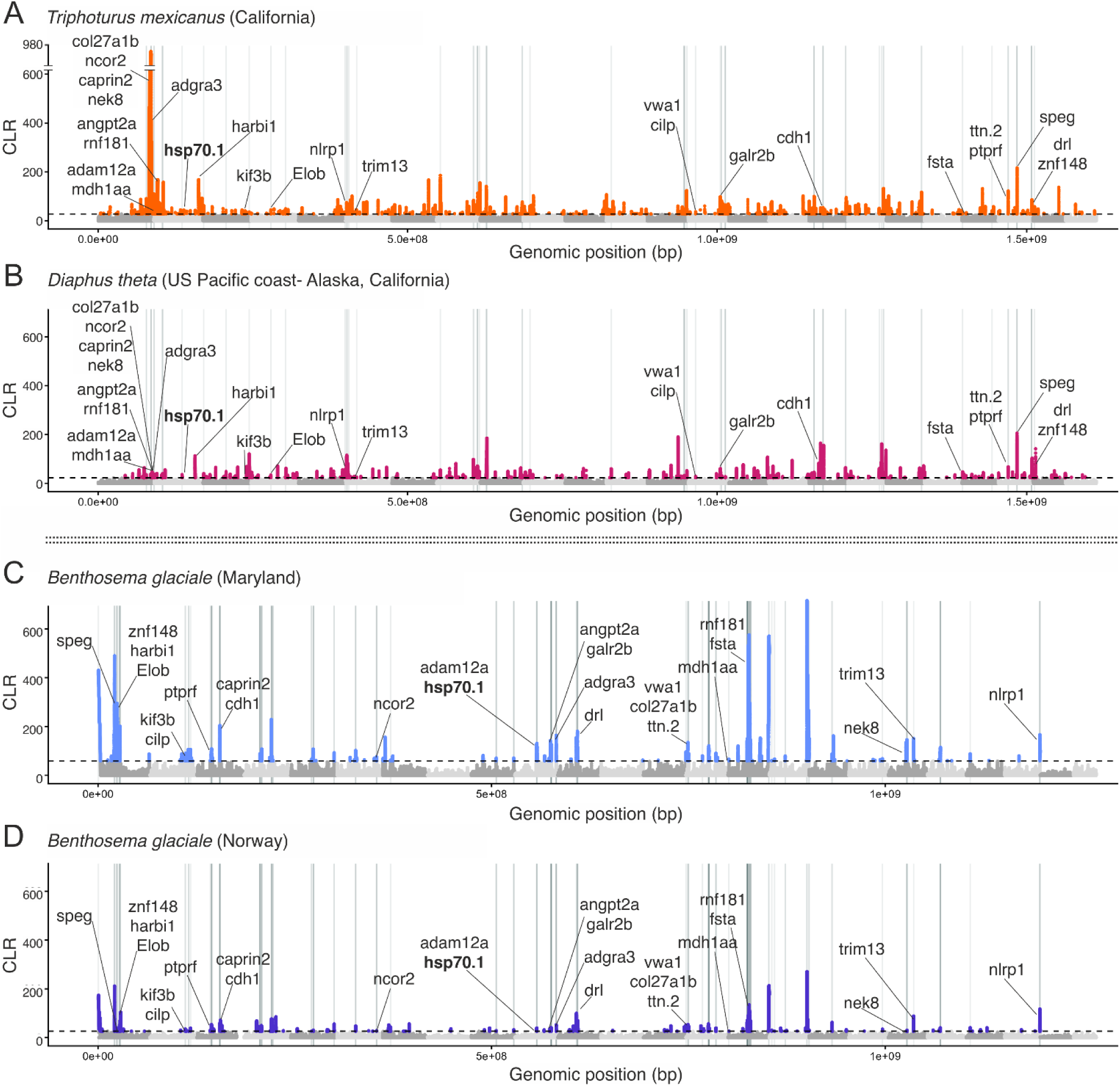
Shared selective sweeps across Atlantic and Pacific lanternfishes. Peaks in composite likelihood ratio (CLR) from SweepFinder2 (Degiorgio et al., 2016) across the two pairs of populations, separated by the double dotted line. Dashed horizontal lines indicate a conservative 2% threshold for detecting selective sweeps; all colored peaks exceeded this threshold, calculated separately for each focal species. Vertical lines indicate shared CLR peaks between A) California *Triphoturus mexicanus* and B) *Diaphus theta* or between C) Maryland *Benthosema glaciale* and D) Norway *B. glaciale.* Labeled candidate genes (Table 2) occurred within 50kb of these CLR peaks in all four datasets. The gene *hsp70.1,* a major candidate in climate change adaptation, is shown in bold across all four populations. Chromosomes are represented by alternating colors. Note that *B. glaciale* and *Triphoturus/Diaphus* were aligned to different reference genomes, so the x-axes do not align across each species pair. For exact gene locations, see Supp. Table 3-4.

Within these coding regions under selection, 313 out of 523 *B. glaciale* and 145 out of 543 *Triphoturus/Diaphus* protein sequences produced high quality matches (E<10^-5^) with the zebrafish Swissprot database. Several sequences shared gene identity, so that the *B. glaciale* matches added up to 255 distinct genes, and the *Triphoturus/Diaphus* matches added up to 59 distinct genes.

### Enriched gene ontologies across oceanic basins

Ontology was assigned to candidate genes based on the zebrafish gene ontology. 203 GO terms were found to be significantly enriched in the Atlantic dataset and 17 in the Pacific dataset (P < 0.01), with seven GO terms being significantly overrepresented in both Atlantic and Pacific lanternfishes (Fig. 4). Six out of these seven shared ontologies were related to sensory organ development, particularly the otolith/inner ear. Retina development was the only term not related to ear structures. The seventh shared GO term was dorsal/ventral axis pattern formation (Fig. 4). In the Pacific, there was complete overlap between genes in this category and genes in the ear development categories, whereas the Atlantic included seven genes not found in the inner ear development group. Overlap between enriched GO terms in the two oceanic basins does not imply overlap in the identity of genes involved in the significant ontologies: while most shared enriched terms are related to ear morphogenesis, only two ear morphogenesis genes (*cdh* and *kif3b*) share signals of selection across all populations and oceanic basins (Table 2).

**Figure 4.**
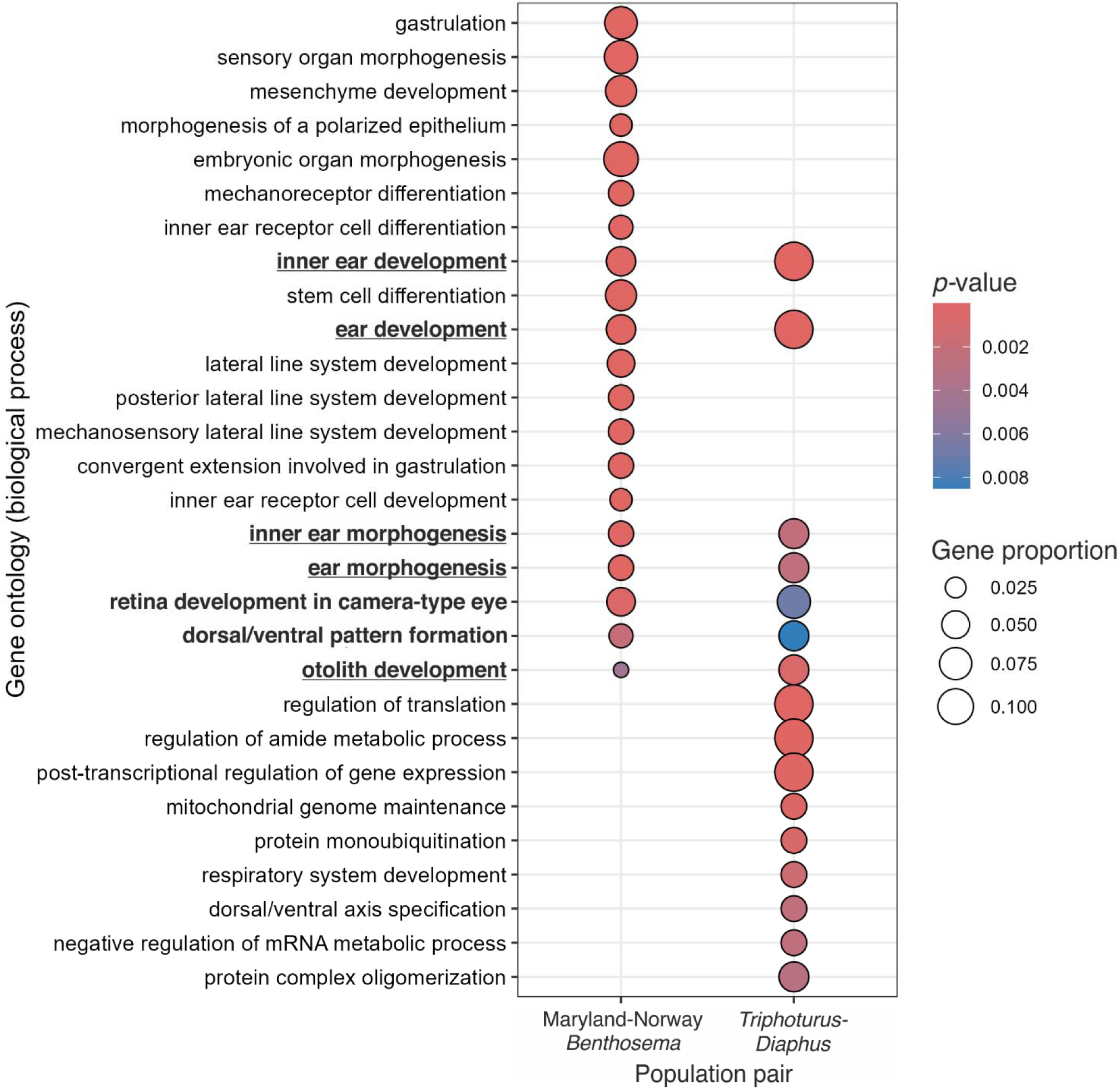
Significantly enriched gene ontology (GO) terms among genes under likely selection in Atlantic (Maryland/Norway *Benthosema*) and Pacific (*Triphoturus/Diaphus*) lanternfishes. The categories shown represent all seven ontologies enriched in both oceanic basins (highlighted in bold) and the top 15 most significantly enriched ontologies in each individual oceanic basin. Shared terms strictly related to ear structures are underlined. Color represents the *p-*value and size of the circles indicates the proportion of genes in each dataset associated with each GO term.

### Shared candidate genes under recent selection in all taxa

We identified 34 shared candidate genes found in or near selective sweeps in both Pacific and Atlantic lanternfishes (Table 2, Figure 3, Supp. Table 3-4): 12 from direct four-way matching of UniProt identities and 21 from reciprocal BLAST. Some of these genes appear repeatedly across the *Benthosema* and *Triphoturus/Diaphus* 2-way shared sweeps; they were distributed across 45 annotated genes in *Benthosema* and 55 in *Triphoturus/Diaphus.* Repetition of gene identity was observed most commonly in *Triphoturus/Diaphus* transcripts whose orthology was discovered via reciprocal BLASTp against the *Benthosema* annotation.

We discovered that 79% (27) of the shared candidate genes had functions that could be attributed to at least one relevant climate stressor, including thermal stress response, response to hypoxia, response to acid stress (high CO2 concentrations), biomineralization, Ca^2+^ binding/modulation, or otolith development (Table 2). 14 (42%) of the 34 shared candidate genes were identified as candidate genes in past experimental climate change research on responses to ocean acidification (Table 2): 12 directly and two via close orthologs. Only seven out of 34 genes lacked any direct relevance or putative connection to OA or warming (Table 2).

### Shared candidate gene functions

#### Thermal stress

8/34 candidate genes (Table 2) are directly involved in response to thermal stress, including heat shock response. One gene, *hsp70.1,* encodes the heat shock protein (HSP70). Seven additional genes were involved in heat stress response: *adgra3* (Himanen et al., 2022), *elob* (elongin B) (Y. Liu et al., 2022), *mdh1aa* (Vicentini et al., 2024)*, nlrp1* (Meng et al., 2025)*, ptprf* (Ramírez-Calero et al., 2023)*, rnf181* (Díaz-Díaz et al., 2020; ZFIN Staff, 2022), and *trim13* (George et al., 2023). 6/8 of these genes are implicated in experimental literature on response to heatwaves or thermal stress specifically in the context of climate change, with *hsp70.1* receiving the most attention (Clark et al., 2008; Jeyachandran et al., 2023; G. C. Li et al., 1992; Y. Liu et al., 2022; Suo et al., 2022; Tripp-Valdez et al., 2019; Urbarova et al., 2019; Xu et al., 2022; Yusof et al., 2022; X. Zhang et al., 2022). For the two remaining genes, *rnf181* is responsible for protein ubiquitination and *adgra3*is activated by heat shock factors (Himanen et al., 2022).

#### Ocean acidification

10/34 candidate genes are involved in acid stress response. Organismal and cellular response to acid stress has been extensively tested in the context of OA, and all of these genes’ products are well-featured in the relevant literature: *adam12a* (Ertl et al., 2016; Moya et al., 2016)*, angpt2a* (Maas et al., 2024)*, cdh1* (Downey-Wall et al., 2020)*, cilp* (Moya et al., 2016)*, col27a1b* (Mittermayer et al., 2019)*, fsta* (Ertl et al., 2016)*, harbi1* (Spencer et al., 2026)*, hsp70.1* (Urbarova et al., 2019)*, ttn.2* (Artigaud et al., 2015; Goncalves et al., 2017; Spencer et al., 2026)*, ptprf* (Ramírez-Calero et al., 2023), and *nlrp1* (Meng et al., 2025; Y. C. Wang et al., 2015). F*sta* is also involved in other forms of stress response (Kuganathan et al., 2026). All genes in this category are linked to organismal responses to reduced pH caused by CO2 accumulation, making this category synonymous with CO2 stress.

#### Hypoxia

10/34 candidate genes are linked to hypoxia either directly (3/10) or through their role in hemopoiesis (7/10), heart and blood vessel development, or the circulatory system in general. The circulatory system genes *angpt2a* (Abaci et al., 2010; Gustavsson et al., 2007; Huth & Place, 2016), *drl* (Robertson et al., 2016), *ttn.2* (Krishnan et al., 2008; Zhou et al., 2020), *ncor2* (Arango et al., 2026; Tijssen et al., 2011), *nek8* (Manning et al., 2013), *speg* (Quick et al., 2017) and *znf148* (X. Li et al., 2006; Woo et al., 2008) are joined by *adam12a* (also involved in biomineralization and response to acid stress)(Allen et al., 2020), and the aforementioned heat shock response genes *adgra3* (Himanen et al., 2022) and *hsp70.1* (Suo et al., 2022; Tripp-Valdez et al., 2019). The three genes that have been featured in climate change literature for their activity under hypoxic stress are *angpt2a* (Huth & Place, 2016)*, ttn.2* (Zhou et al., 2020) and *hsp70.1* (Suo et al., 2022; Tripp-Valdez et al., 2019), which are also known to have a role in other forms of OA/climate change stress responses (Table 2).

#### Biomineralization

8/34 candidate genes are involved in biomineralization. Of these, *col27a1b* (Christiansen et al., 2009; Hjorten et al., 2007) and *fsta* (Gajos-Michniewicz et al., 2010; Sylva et al., 2013) are directly related to skeletal system development (Bradford et al., 2022). Other skeletal system genes in our list are *cdh1* (Chandra Rajan et al., 2021), *cilp* (Johnson et al., 2003; Mori et al., 2006) and *vwa1* (Chandra Rajan et al., 2021; Niu et al., 2023), involved in cartilage development, and *kif3b* (Ning et al., 2026) and *adam12a* (Atfi et al., 2007; Ren et al., 2017; Tokumasu et al., 2016), implicated in osteoarthritis. The last gene in this category is *galr2b* (galanin receptor 2b), recently found to have a novel role in craniofacial development through functional studies of embryonic jaw development in San Salvador island pupfishes (Palominos et al., 2023). A*dam12a, cdh1, cilp, col27a1b, fsta* and *vwa1* (6/8) have all been featured in experimental studies of responses to climate change, though *vwa1* (Chandra Rajan et al., 2021; Vargas et al., 2022; Yuan et al., 2025) is the only one of these to have been discussed specifically in the context of biomineralization.

#### Calcium modulation

8/34 candidate genes either bind calcium, are modulated/activated by calcium, or regulate calcium intake and homeostasis. Theoretically this category has partial overlap with biomineralization (discussed above), but it is meant to encompass any genes that may act upstream of biomineralization (Di Santo, 2019), in Ca^2+^ transport and management. Of these nine, *caprin2* is the only gene to have no other known relevant functions to climate change (Miao et al., 2014) whereas other genes in this list overlap with other relevant functions: *adam12a* (Nyren-Erickson et al., 2013)*, cdh1* (Chandra Rajan et al., 2021)*, kif3b* (Phang et al., 2014)*, ttn.2* (Labeit et al., 2003)*, nek8* (Roig, 2025)*, speg* (Campbell et al., 2021), and *vwa1* (Ambort et al., 2012).

#### Otolith development

Two out of 34 candidate genes relate to inner ear morphogenesis and otolith formation: *cdh1,* implicated in a study on the effect of ocean acidification on balance in the yellow croaker (X. Wang et al., 2023) and *kif3b* (Zhao et al., 2012).

## Discussion

We present evidence of shared signals of selection on 34 candidate genes across three lanternfish species in two oceanic basins, many of which are functionally linked to environmental stressors associated with climate change, including thermal stress, hypoxia, acidification, and biomineralization. The candidate genes and shared ontology terms paint a multidimensional and global picture of organismal response to recent selective pressures, including genes involved in oxygen transport, acidic pH tolerance, heat-shock response, and biomineralization. We did not detect any recent declines in effective population size, but we do find evidence of historical population expansion post-glaciation.

### Population size and historical population expansion

For all three lanternfish species, we estimated a near-constant effective population size (N_e_) of 5 million individuals over the past 200 generations/100 years, suggesting that the effective number of individuals contributing offspring in each generation has remained stable during anthropogenic climate change. However, this estimate is likely far lower than the census population size of these species and may result from multiple factors. First, like many marine fishes, myctophiforms employ a sweepstakes reproductive strategy: despite each individual producing thousands of eggs, most offspring die before reaching reproductive maturity and reproductive success is highly skewed. This is known to cause extreme N_e_/N_c_ ratios of 10^-3^ or less (Hedgecock & Pudovkin, 2011; Waples, 2016), not uncommon in abundant marine fishes (Feng et al., 2017; Marandel et al., 2019; Waples, 2016), and also supported by recent RADseq estimates of genetic diversity for three Myctophidae species in the Gulf of Mexico (Bernard et al., 2022). Second, selective sweeps within the population, as we detected in this study, are known to effectively depress N_e_ by reducing genetic diversity in localized regions of the genome (Buffalo, 2021; Charlesworth & Jensen, 2021; Comeron et al., 2008; Ellegren & Galtier, 2016). One recent study of *Benthosema glaciale* mitochondrial DNA in the Mediterranean Sea estimated a comparable N_e_ of ten million individuals, with large posterior density intervals overlapping our estimate of 5 million (Sarropoulou et al., 2022).

Across all three species and oceanic basins, Tajima’s D was strongly negative, indicating a recent population expansion and/or strong purifying and positive selection, and genetic diversity (π) was lower than predicted (0.0015-0.0023) . However, in agreement with Lewontin’s paradox, these values of π are not unexpected for extremely abundant pelagic marine organisms (e.g. Atlantic herring, π=0.003 (Feng et al., 2017; Mohamadnejad Sangdehi et al., 2024) and phytoplankton *Emiliania huxleyi* π=0.006 (Filatov, 2019)) due to efficient purging of mutations and an abundance of selective sweeps (Filatov, 2019). Accordingly, we find that in our sampled populations, π is lower in regions identified as likely selective sweeps. Additionally, in the Atlantic herring, low π and N_e_ of 400,000 are attributed to low mutation rate and recent expansion from a historical population bottleneck (Feng et al., 2017). While the mutation rate is not known for any myctophiform, it is plausible that their low π and estimated N_e_ may similarly be due to the same factors combined with the effect of sweeps on genetic diversity. Given their large population sizes, a historical myctophiform bottleneck may date back to the last glacial maximum (LGM), which affected multiple pelagic fish species (Shum et al., 2015; Silva et al., 2014) and all but one (San Diego) of our sampling locations. The least affected species in this scenario would be *T. mexicanus* with its exclusively sub-Arctic distribution, which indeed shows the highest π and least negative Tajima’s D in four populations.

### Shared gene ontology across species and oceanic basins

Most importantly, our results reveal striking shared trends in recently selected genes and ontologies across different, distantly related myctophid species collected in disparate locations and environments (Fig. 3-4, Table 2). Many sweeps were shared between *B. glaciale* from Norway and Maryland and between California *T. mexicanus* and *D. theta* from California and Alaska (Fig. 3).

We found shared enriched GO terms under selection in the Atlantic and Pacific for inner ear morphogenesis, including otolith formation (Fig. 4). One of the most well-documented effects of OA on fish physiology is abnormal otolith development (Bignami et al., 2013; Holmberg et al., 2019; Pimentel et al., 2014; Wexler et al., 2023), a phenomenon that has been extensively tested in lab environments and observed in disparate fish families (Bignami et al., 2013; Holmberg et al., 2019; Pimentel et al., 2014; Wexler et al., 2023). Otoliths (ear stones) are impacted by alterations to the biomineralization process in more acidic water, analogous to invertebrate shells and vertebrate skeletons, yet they also show phenotypic plasticity in response to suboptimal environmental conditions (Schulz-Mirbach et al., 2019). A more acidic environment produces overgrown otoliths in most tested species (reviewed in (Holmberg et al., 2019)), a reaction that has implications for fish sensory perception, movement and behaviour (Ashur et al., 2017). The positively selected inner ear genes we observe here in Myctophidae may indicate rapid adaptation to increasing ocean acidification over the past half century. This study adds to the mounting evidence of otoliths as a major trait of interest in fish responses to OA, and the first time this evidence has been documented in mesopelagic fishes at a global scale.

The GO term for retina development in camera-type eyes (Ashur et al., 2017) is likewise involved in sensory systems. Exposure to high CO2 concentrations alters fishes’ response to light, causing hypersensitivity and impaired anti-predatory response (Ashur et al., 2017), but it is unknown how this may affect mesopelagic fishes given their unusual eye developmental processes and adaptation to a low-light environment (Fogg et al., 2024). It may impact survival during the lanternfishes’ larval stage, when they reside in the more light-rich epipelagic zone (0-200m) (Sassa, 2019; Sassa et al., 2015). The last enriched GO term was dorsal/ventral pattern formation, a process that is regulated by BMP (Bone Morphogenetic Protein) signalling (Bier & De Robertis, 2015; Pomreinke et al., 2017) and may be connected to biomineralization.

### Shared gene identity across species and oceanic basins

We discovered that a surprising majority of the 34 candidate genes with strong signals of positive selection in all three species appear to be directly involved in responses to OA and climate change. This list is not comprehensive: many genes located within or in proximity to selective sweep regions remain unknown due to lack of annotation, and our methods are geared towards detection of hard sweeps, potentially leaving any soft sweeps unexplored. We also have not shown a direct functional link between any of the 34 candidate genes and selective pressures from OA/climate change. However, it was possible to assign 27/34 genes to one or more relevant physiological pathways known to be involved in adaptation to climate change in other taxa (Table 2). Many of these physiological functions are connected to each other: biomineralization, Ca^2+^ modulation, and otolith formation are linked through their dependence on calcium; thermal stress and oxygen demand are linked through the fish’s cardiovascular response to high temperatures (Farrell et al., 2009); the circulatory system itself is modulated by Ca^2+^ homeostasis, connecting hypoxia to Ca^2+^ modulation (Tibbits et al., 1991). In summary, adaptation to a more acidic, less oxygen-rich, warmer ocean clearly involves multiple interconnected physiological processes. Most of the 34 genes were involved in more than one of these physiological processes, notably *hsp70.1* (Y. Liu et al., 2022; Suo et al., 2022; Tripp-Valdez et al., 2019; Urbarova et al., 2019; Xu et al., 2022; Yusof et al., 2022)*, cdh1* (Chandra Rajan et al., 2021; Downey-Wall et al., 2020; X. Wang et al., 2023) and *ttn.2* (Artigaud et al., 2015; Goncalves et al., 2017; Ryu et al., 2020; Spencer et al., 2026; Zhou et al., 2020), well-known genes involved in a wide range of stress responses to experimental studies of climate change stressors (Table 2).

### Stress response candidate genes

Heat shock proteins (HSPs) were of primary interest due to their involvement in the cytological stress response (Hu et al., 2022). Climate change and OA subject organisms to various stressors: hypoxia, increased water temperatures, longer heatwaves, issues maintaining proper ion homeostasis. While HSPs were initially named after their activation in thermal stress, any of these phenomena may trigger their expression. One of the shared genes under selection, *hsp70.1,* is a major player in adaptation to thermal stress and of great interest in research on how organisms react to climate change (Jeyachandran et al., 2023). Many studies in the past have highlighted HSP70’s potential role in tolerance to climate change in marine ecosystems, not only in response to heat stress (Y. Liu et al., 2022; Xu et al., 2022; Yusof et al., 2022), but also to hypoxia (Suo et al., 2022; Tripp-Valdez et al., 2019) and elevated CO2 concentrations (Urbarova et al., 2019). HSPs are also involved in thermal stress response to diel migration in crustaceans (Bernatowicz et al., 2021; Elder & Seibel, 2015). Higher levels of HSP70 expression are linked to increased behavioural plasticity in *Daphnia* under thermal stress, which allows them to better cope with sudden environmental changes (Bernatowicz et al., 2021). A similar role may extend to lanternfishes during diel migrations.

HSP70 is directly linked to two more genes under selection, *EloB* and *adgra3.* HSP70’s expression is regulated by the elongin complex, including *EloB’*s product Elongin B (Gerber et al., 2005; Kawauchi et al., 2013; Sheng et al., 2024). *EloB* was identified as a central gene in thermal stress response in the Pacific oyster *Crassostrea gigas,* where it works alongside HSP70 (Y. Liu et al., 2022). HSP70 is also connected to *adgra3’*s via their induction by heat shock factors (HSF) (Himanen et al., 2022; Joutsen et al., 2020). *Adgra3* expression is induced by HSF2 under conditions of oxidative and thermal stress (Himanen et al., 2022) and it encodes one of many adhesion proteins targeted by HSFs, as cell-cell adhesion is important for cellular survival to heat damage (Himanen et al., 2022; Joutsen et al., 2020). We discovered additional candidate genes (Table 2) that respond to heat stress, ranging from essential metabolism to protein ubiquination (*rnf181, trim13*), to neurophysiological (*ptprf*) and innate immune responses (*nlrp1*).

### Biomineralization candidate genes

Biomineralization is widely impacted by ocean acidification in marine invertebrates (Armstrong & Bahr, 2025; Hofmann & Schellnhuber, 2009; Maas et al., 2024; Pörtner, 2008). In the context of vertebrate physiology, this translates to a projected effect on bone development that has already been demonstrated in some species (Di Santo, 2019; Pimentel et al., 2014; Sundin, 2023). Thus, bone development/biomineralization genes were one of the primary categories of interest for this analysis. Two of the 34 genes under selection are directly related to skeletal development as per their gene ontology: *col27a* and *fsta* (Bradford et al., 2022; Christiansen et al., 2009; Hjorten et al., 2007). Collagens like *col27a*’s product are involved in response to OA in corals living in volcanic CO2 seeps (Leiva et al., 2023). Meanwhile, *fsta’*s product follistatin-A is a crucial regulator of bone development and mineralization (Gajos-Michniewicz et al., 2010; Graf et al., 2016). Experimental evidence for a role of *fsta* in response to OA has been found in at least one marine oyster, where it is strongly upregulated at elevated CO2 (Ertl et al., 2016). It is overall both a skeletal development protein and a stress response protein (Kuganathan et al., 2026; L. Zhang et al., 2018).

Several more genes are broadly involved with biomineralization even while lacking bone morphogenesis as an ontology term (Table 2). For example, *vwa1* appears in several papers on the effect of climate change for its role in biomineralization (Vargas et al., 2022; Yuan et al., 2025; T. Zhu et al., 2023), but its primary function is upstream in cartilage conformation (Niu et al., 2023).

Compared to the abundance of existing scientific literature on invertebrate calcification pathways in OA (Armstrong & Bahr, 2025; Chandra Rajan et al., 2021; Di Santo, 2019; Maas et al., 2024; Vargas et al., 2022), the vertebrate skeletal system’s response to OA in natural populations is still not fully understood, making it difficult to pinpoint how selection is affecting their function. With many of the genes under selection being involved in the TGFβ-BMP pathways, in control of cartilage/bone localization, or having mutants implicated in osteoporosis/osteoarthritis, it is likely that their role is mostly in preserving skeletal integrity in stressful conditions.

### Hypoxia, calcium, and otolith candidate genes

OA diminishes oxygen availability (Gobler & Baumann, 2016; Hofmann & Schellnhuber, 2009; Venegas et al., 2023) so widespread adaptations of cardiac tissues or blood that improve survival in hypoxic environments may be expected. This is reflected by the large proportion of candidate genes active in the circulatory system. These include *ttn.2*, encoding the giant protein titin responsible for heart contraction and myofibril assembly (Bradford et al., 2022), whose expression has been found in past research to be impacted by all major climate change stressors (Table 2). Titin is functionally linked to *speg* (striated muscle preferentially expressed protein kinase), a key regulator of cardiac calcium homeostasis (Campbell et al., 2021; Quick et al., 2017), connecting oxygen circulation with another key aspect of OA, the decrease in calcium availability.

Experimental treatments that exposed marine invertebrates to acidified waters showed an upregulation of genes responsible for calcium management (X. Wang et al., 2020; Xin et al., 2022). While Ca^2+^ is naturally relevant to OA in the form of biomineralization, it is a major signaller within cells and important in maintaining homeostasis both within cells and in the extracellular matrix (Bootman & Bultynck, 2020). Many cardiac system proteins in our list are connected to both hypoxia and Ca^2+^ modulation (Table 2), including Speg (Gautel, 2011; Quan et al., 2019). Ca^2+^ is also important in cellular adhesion, another property that has been found to be jeopardized by OA, with most evidence found in sessile invertebrates (Drake et al., 2018; Guan et al., 2025; Kaniewska et al., 2012). We found the gene *cdh1*, encoding the calcium-dependent adhesion protein cadherin-1 (Cailliez & Lavery, 2005; Van Roy & Berx, 2008), to be under likely selection across all species. *Cdh1’*s expression is modulated in response to thermal stress (Guan et al., 2025) and acid stress (Downey-Wall et al., 2020), but cadherins also have a role in biomineralization (Chandra Rajan et al., 2021). *Cdh1* is expressed in the fish inner ear and is involved in otolith morphogenesis, with high CO2 concentrations likely disrupting this function (X. Wang et al., 2023). This makes *cdh1* one of two otolith genes under shared selection across all species/oceanic basins, the other being *kif3b* (Ning et al., 2026; Whitfield, 2020; Zhao et al., 2012), also fine-tuned by calcium (Phang et al., 2014).

### Lanternfish ecology and adaptation to climate change

Myctophiform and deep sea ecology must be considered when interpreting the findings described above. The effects of climate change, including both global warming and OA, partly decrease with depth, particularly below 700 m (IPCC, 2013; Lindsey & Dahlman, 2025; NOAA, 2020). Compared to coastal species, lanternfishes may experience less severe environmental pressures from climate change. However, two aspects of their life history increase their vulnerability. 1) Like many mesopelagic species, myctophid eggs and larvae develop in the epipelagic zone (Sassa, 2019; Sassa et al., 2014, 2015). Larval fishes are usually more vulnerable to habitat fluctuations than adults (Pankhurst & Munday, 2011), consequently most climate-related mortality and the harshest abiotic selective pressures may occur during this life stage. 2) Most myctophids perform diel migration to the epipelagic zone to feed at night (Catul et al., 2011; Knutsen et al., 2023), and it is during these migrations that they are likely to be exposed to the more extreme surface conditions, with behavioural and physiological effects. Vertically migrating taxa in general show a stronger response to climate change than non-migrating taxa (Hsieh et al., 2009; Seibel & Birk, 2022).

Our results from demographic and selection analyses suggest that lanternfishes are not decreasing in overall abundance in response to recent anthropogenic climate change, but are adapting their physiology to new global environmental pressures from OA and warming through the involvement of HSP70 and other proteins that may promote cellular repair, resilience, and homeostasis (Table 2) (Jeyachandran et al., 2023). Physiological adaptations may in turn come with behavioural changes, which have already been documented in some cases. For example, following a severe heatwave in 2015-2016 in California, local myctophid communities adjusted their foraging preferences to the deeper ocean, which likely impacted local carbon transport and food webs (Iglesias et al., 2024). As the severity of heatwaves increases, these behavioural shifts may be observed more frequently and across more locations, affecting the global carbon cycle (Iglesias et al., 2024). At the same time, future scenarios of global warming may lead to range shifts, including range expansion towards higher latitudes in lanternfish taxa currently found in tropical waters as predicted for other vertically migrating taxa (Seibel & Birk, 2022; Freer et al., 2019).

## Conclusion

Myctophiformes are keystone taxa across all oceans and a central component of the global carbon cycle. Yet we previously had no knowledge of their responses to the accelerating impacts of anthropogenic climate change. We used population genomics to identify genes responding to shared environmental selective pressures in parallel across species spanning the Atlantic and Pacific oceans and discovered that a large majority of these shared candidate genes have potential functions in adaptation to OA and thermal stress resilience, well-supported by experimental literature on the topic. This research provides a new framework for studying global adaptations to climate change across taxa and offers unprecedented insight into the global shared responses of hyperabundant mesopelagic vertebrates to the ongoing effects of climate change.

## Acknowledgments

This research was funded by NSF DEB 2519904 and NIH 5R01DE027052-02 awards to CHM. We thank the RACE Groundfish and Shellfish Assessment Programs of the NOAA Fisheries Alaska Fisheries Science Center and the crew of the R/V NorthWest Explorer and R/V Alaska Knight (Eastern Bering Sea Groundfish Bottom Trawl Survey), the Scripps Institution of Oceanography and the crew of the R/V R.G. Sproul for their assistance in securing Pacific Ocean samples. We thank the crew onboard R/V G.O. Sars and Francesco Saltalamacchia for securing fish samples from Fensfjorden as part of a survey funded by the University of Bergen and The Research Council of Norway (HypOnFjordFish; Project number: 301077). We thank Alex Davis for generously providing Baltimore Canyon specimens of *B. glaciale*. We thank Shaoxiong Ding (Xiamen University) for providing us with up-to-date annotation of the *B. pterotum* reference genome. We also thank Rebecca Abrams, Ichthyology collection manager at the Museum of Vertebrate Zoology, UC Berkeley, for cataloguing all samples used in this study (MVZ:Fishes:XX). All research procedures, euthanasia methods, and animal care protocols (AUP-2021-02-14062-1 and AUP-2021-07-14515) were approved by the University of California, Berkeley Animal Care and Use committee. OpenAI’s ChatGPT was used exclusively for assistance with bash scripts and R coding.

---

**Supplementary Figure 1.**
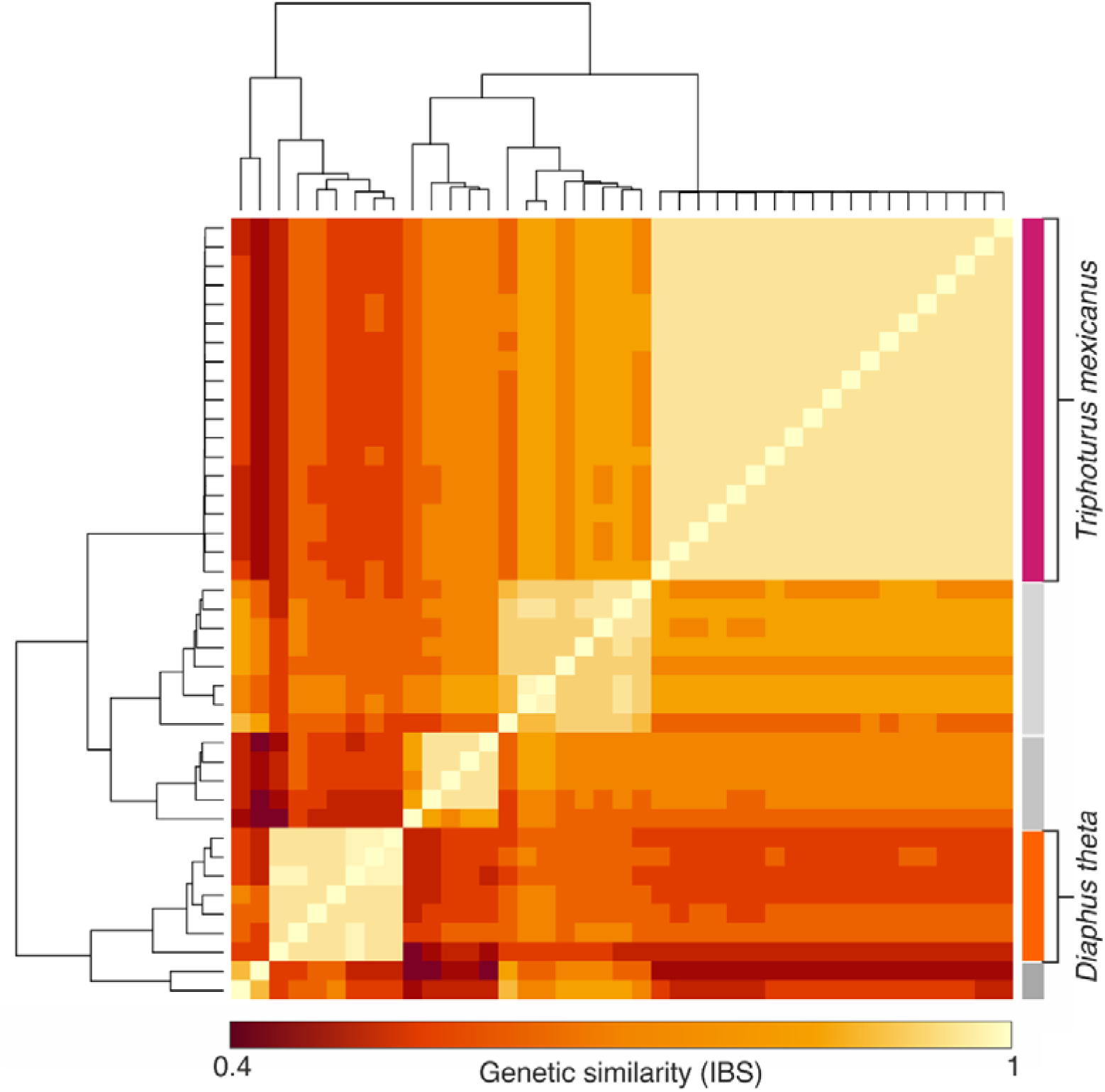
Clustering of samples based on genomic similarity in the Lampanyctini and Diaphini clades across a set of independent SNPs, pruned for linkage disequilibrium using PLINK 1.9 (Purcell et al., 2007). This analysis was run without *a priori* awareness of species identity. Genetic similarity is given as identity-by-state (IBS), representing the proportion of shared alleles (no two samples shared less than 40% of alleles). Of the recovered clusters, two were used for downstream analyses based on robustness and number of specimens, corresponding to individuals later identified morphologically as *Triphoturus mexicanus* and *Diaphus theta*.

**Supplementary Figure 2.**
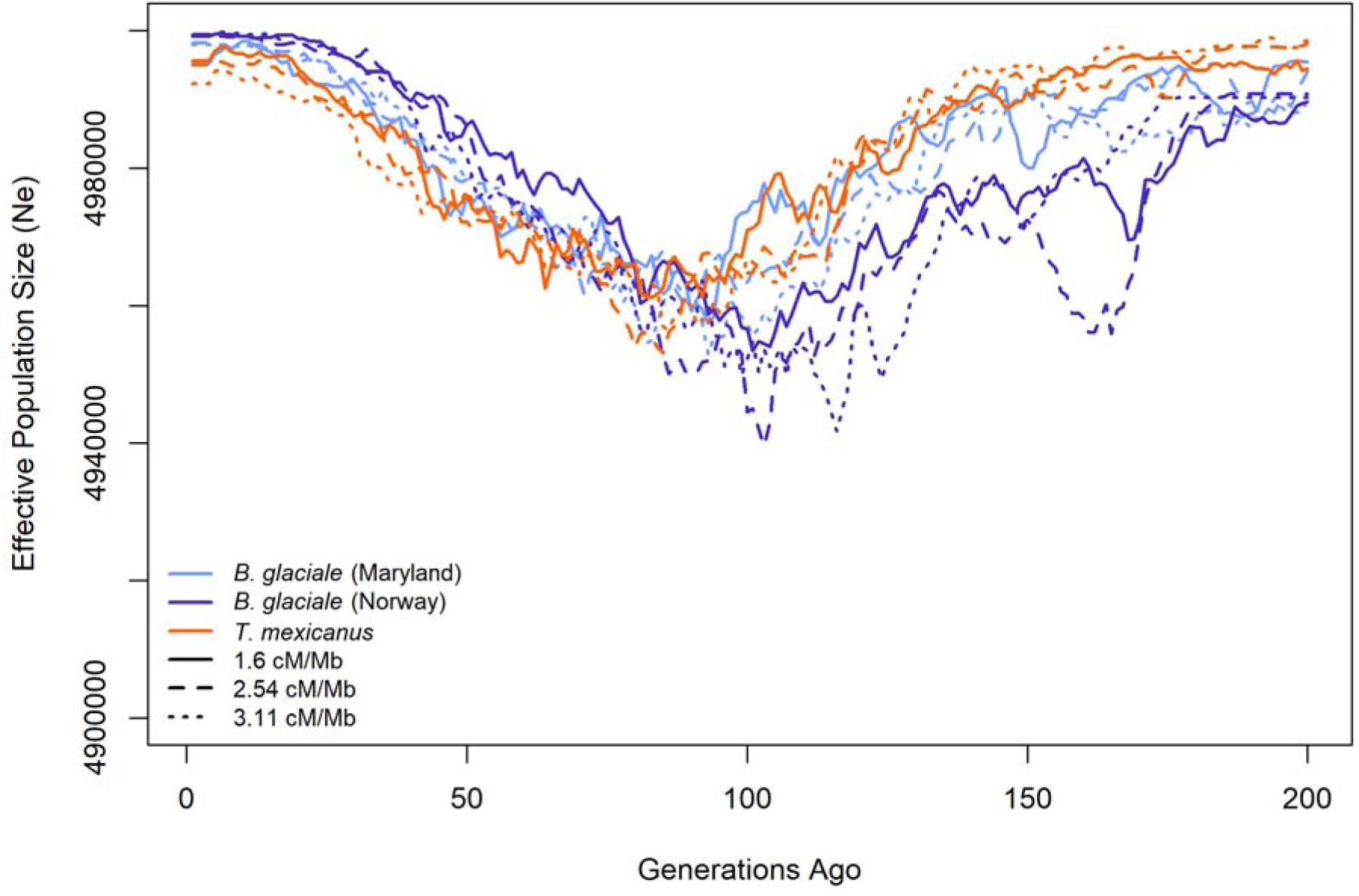
Effective population size (N_e_) as estimated by GONe (Santiago et al., 2025) over the past 200 generations in *Benthosema glaciale* from Norway and Maryland, and in *Triphoturus mexicanus.* As recombination maps are not available for myctophids, three different recombination rates were tested (assumed constant): 1.6 cM/Mb from the zebrafish (Bradley et al., 2011); 2.54 cM/Mb from the Atlantic herring (Pettersson et al., 2019); and 3.11 from the threespine stickleback (Wang et al., 2026). A minor fluctuation in N_e_ at -100 generations, consistent across all tests, is emphasized by the y-axis scale, but is likely not biologically meaningful given the magnitude of N_e_ (Gargiulo et al., 2024).

